# Intravenous whole blood transfusion results in faster recovery of vascular integrity and increased survival in experimental cerebral malaria

**DOI:** 10.1101/2021.12.26.474205

**Authors:** Saba Gul, Hans C. Ackerman, Cláudio Tadeu Daniel-Ribeiro, Leonardo J. M. Carvalho

## Abstract

Transfusion of 10 mg/kg of whole blood via intraperitoneal route to mice with late-stage experimental cerebral malaria (ECM) along with artemether has been shown to result in markedly increased survival (75%) compared to artemether alone (51%). Intraperitoneal route was used to overcome the restrictions imposed by injection of large volumes of viscous fluid in small and deranged blood vessels of mice with ECM. In the present study, a method of intravenous transfusion was implemented by injecting 200μL of whole blood through the right jugular vein in mice with late-stage ECM, together with artemether given intraperitoneally, leading to a remarkable increase in survival, from 54% to 90%. On the contrary, mice receiving artemether plus plasma transfusion showed a worse outcome, with only 18% survival. Compared to the intraperitoneal route, intravascular transfusion led to faster and more pronounced recoveries of hematocrit, platelet counts, angiopoietins levels (ANG-1, ANG-2 and ANG-2/ANG-1) and blood brain barrier integrity. These findings indicate that whole blood transfusion when given intravenously show more efficacy over intraperitoneal transfusion, reinforcing evidence for benefit as an adjuvant therapy for cerebral malaria.

## Introduction

Despite advances in medical science and global technical strategies for malaria control, there were an estimated 229 million malaria cases with 409,000 deaths in 2019 [1]. Cerebral malaria (CM) is prominently involved in majority of malaria related deaths, and frequently results in long-term cognitive deficits in survivors [2,3]. Majority of deaths befall in the first 24-48 hours of hospital admission, demanding a quick administration of treatments to reduce death [4,5]. In order to address these issues, to increase survival and overcome the occurrence of post treatment neurological sequelae, there is utmost need of adjuvant therapies.

Cerebral malaria is a multifactorial disorder accompanied by hematological and vascular changes, including mild to moderate to severe anemia, altered hemorheologic properties, thrombocytopenia, along with alterations of biochemical characteristics of the plasma such as the levels of angiopoietins, haptoglobin, L-arginine and proinflammatory cytokines, among others [6–12]. Recently, a multicenter study in Africa showed that children hospitalized with severe *P. falciparum* malaria, particularly those with impaired consciousness or severe acidosis, benefited from whole blood transfusion, exhibiting increased survival [13,14]. Accordingly, whole blood transfusion, when administered intraperitoneally to mice with late-stage ECM also receiving artemether, resulted in increased survival and had beneficiary effects in terms of restoring hematocrit and markers of vascular health [15]. These findings with whole blood transfusion in ECM follow prior evidence indicating that interventions that restore vascular function are beneficial as adjunctive therapy in ECM [16–19].

In the study showing the effect of blood transfusion along with artemether in ECM, transfusion was performed by means of intraperitoneal injection. This route was chosen for its simplicity and due to the complicated task of injecting relatively large amounts (200μL) of a viscous fluid in the commonly used lateral tail vein in very sick mice with ECM, which shows vasoconstriction and vascular plugging by adherent monocytes [16,18–23]. The downside of this strategy is that it does not properly mimic the intervention as it happens in humans and, as a consequence, the benefit of transfusion may not be as immediate as it would be with an intravenous injection. Besides, not 100% of the injected red blood cells reach the circulation [24] and it is expected that losses occur as well for the transfused plasma components. Therefore, in the present study we performed whole blood transfusion as adjunctive therapy to artemether in mice with late-stage ECM by means of intravenous injection of heparinized blood using a technique of right jugular vein injection. The results show that the outcome in terms of survival, which reached 90%, was markedly improved compared to intraperitoneal transfusion as previously described [15], and recovery of hematological and vascular parameters was also more prominent.

## Materials and methods

### Mice

The Institute of Science and Technology in Biomodels (ICTB) of FIOCRUZ provided specific pathogen-free, female, 8-12-week-old C57BL/6 mice (16-20 g). A state of 12-hour light/dark cycle and a constant temperature of 21–22 °C was maintained where animals had free access to water and chow. In experimental protocols every regulation and guideline were properly followed. All the experimental work has been ratified by Animal Welfare Committee of the FIOCRUZ (CEUA/FIOCRUZ), under license number L-037/21.

### Infection with *Plasmodium berghei* ANKA (PbA)

For passage, healthy C57BL/6 mouse were used to cultivate *Plasmodium berghei* ANKA (PbA) parasites (kindly donated by MR4; the Malaria Research and Reference Reagent Resource Center, Manassas, VA, USA). PbA samples were removed from liquid nitrogen and brought to room temperature and injected to passage mice by intraperitoneal (ip) injection. Infection in experimental group began after intraperitoneal inoculation of 1×10^6^ infected RBC (day 0), and parasitemia assessed on day 6 by flow cytometry or Giemsa-stained blood smears. The onset of cerebral malaria in mice was assessed through clinical signs and body (rectal) temperature, using a thermocouple probe (Oakton^®^ Acorn TM; Oakton Instruments, IL, USA).

### Treatments

In a first set of experiments, on day 6 post inoculation, hypothermic mice (31-36 °C) [25] were randomly allocated in three equal groups to define the effect of treatments on survival rate: *i*) artemether only; *ii*) artemether plus 200 μL of blood and; *iii*) artemether plus 200 μL of plasma. The amount of blood and plasma to be transfused was previously defined [15] (200 μL, approximately equivalent to 10 mL/kg) and were obtained from healthy C57BL/6 mice. Artemether 20mg/ml (Dafra Pharma GmbH, Basel, Switzerland) was injected intraperitoneally at 20 mg/kg [25]. Healthy mice (uninfected) and mice with ECM (untreated) were used as controls. Artemether plus plasma showed the worst outcome in terms of survival, therefore this group was removed from the other experiments (see Results).

### Intravenous blood transfusion and sample collection

All mice received a single intraperitoneal injection of artemether (20 mg/kg), while a group of mice additionally received whole blood into the right jugular vein. Briefly, after anesthetizing mice with 75-100 mg/kg ketamine and 10mg/kg xylazine, mice were kept in ventral recumbent position. Jugular vein was made visible by making small incision on the neck. Blood was transfused using 30G-insulin syringe, the needle was entered into lumen of vein after passing through pectoral muscle. All injections were manually performed and mocking was done to mice receiving only artemether.

Sample collection included collection of whole blood and plasma 6hr and 24hr post treatment. For hematological analysis, whole blood was collected in EDTA-coated microtubes through cardiac puncture. For biochemical analysis, plasma samples were withdrawn from whole blood after centrifuging it at 6,000 rpm for 6 minutes. Later on, plasma was distributed in small aliquots and stored at −80 °C. Same procedure was done to uninfected and ECM mice which served as controls in the study.

### Survival experiments

Experiments were performed to check whether intravenous blood transfusion as adjunctive therapy has a beneficiary effect on survival of ECM mice over only artemether and artemether plus plasma. ECM mice (day 6 of infection, rectal temperature of 31-36 °C) were randomly assigned to three treatments groups: (1) artemether 20 mg/kg ip, (2) artemether 20 mg/kg ip plus 200 μL of whole blood iv and (3) artemether 20 mg/kg ip plus 200 μL of plasma. On daily basis one dose of 20mg/kg artemether ip was administered for 5 consecutive days. After the last dose of artemether, mice were followed for another one week and then euthanized with pentobarbital.

### Study of blood hematological components

At time 0 hour, 6 hours and 24 hours, whole blood samples were collected from controls and treated animals and sent to Institute of Science and Technology in Biomodels (ICTB) of FIOCRUZ for hematological components analysis, using a pocH-100i automated hematology analyzer (Sysmex).

### Enzyme-linked immunosorbent assays (ELISA) for biochemical analysis of plasma components

To determine levels of Angiopoietin 1 & 2, Boster Mouse Angiopoietin 1 & 2 Picokine ELISA kits were used. Prestored plasma samples were brought to room temperature and subjected to 1:5 dilution in PBS and the ELISA performed according to the manufacturer’s instructions. For haptoglobin measurement, plasma samples were diluted 1:5,000 and then Douset ELISA kit was used.

### Assessment of blood-brain barrier permeability

To assess the permeability of the blood-brain barrier at 6 hours and 24 hours after treatment, the animals were anesthetized with urethane (2 mg/g ip) with final volume 150 μL per animal. Each animal received an intravenous injection (in the eye plexus) of 2% Evans blue stain (Sigma) diluted in 1X PBS. After 1 hour of the dye injection, the animals were euthanized and 10 mL of ice-cold saline has been transcardially perfused to them, followed by brain removal. Later the harvested brains were incubated in 3 mL of 99.5% formamide (Sigma) for 48 hours at 37°C. healthy controls and ECM untreated mice went through same procedure. To determine the concentration of the dye leaked out of brain in the formamide, 100μl of formamide from each brain was then separated and the absorbance of each sample was evaluated by spectrophotometry, with a wavelength of 630 nm. A standard curve ranging from 1,285 μg/ml to 1.25 μg/ml was used to calculate the amount of Evans blue extracted per brain tissue.

## Results

### Intravenous blood transfusion prevents artemether-induced anemia

To better mimic a clinically relevant scenario when children present with neurological signs, mice were treated after the development of neurological symptoms on day 6 post infection with artemether (20 mg/kg) alone, artemether plus plasma (200 μL), or artemether plus whole blood (200 μL) IV. Mild to moderate anemia with mean hematocrit value 46.2 % was recorded in ECM mice (uninfected controls: 50.1 %) (**Figure 1A**). The data on hematocrit level per time point showed a continuous decline of anaemia levels (mean: 41.4 % to 33.1 % after 6 hours and 24 hours respectively) in mice treated with artemether alone. On the other hand, treatment with whole blood prevented the further decline in hematocrit (mean: 48.6 % at 6 hours and 46.0 % at 24 hours). Indeed, hematocrit levels in transfused mice at 6 and 24 hours were not different from that observed in ECM mice before treatment.

**Figure 1.**
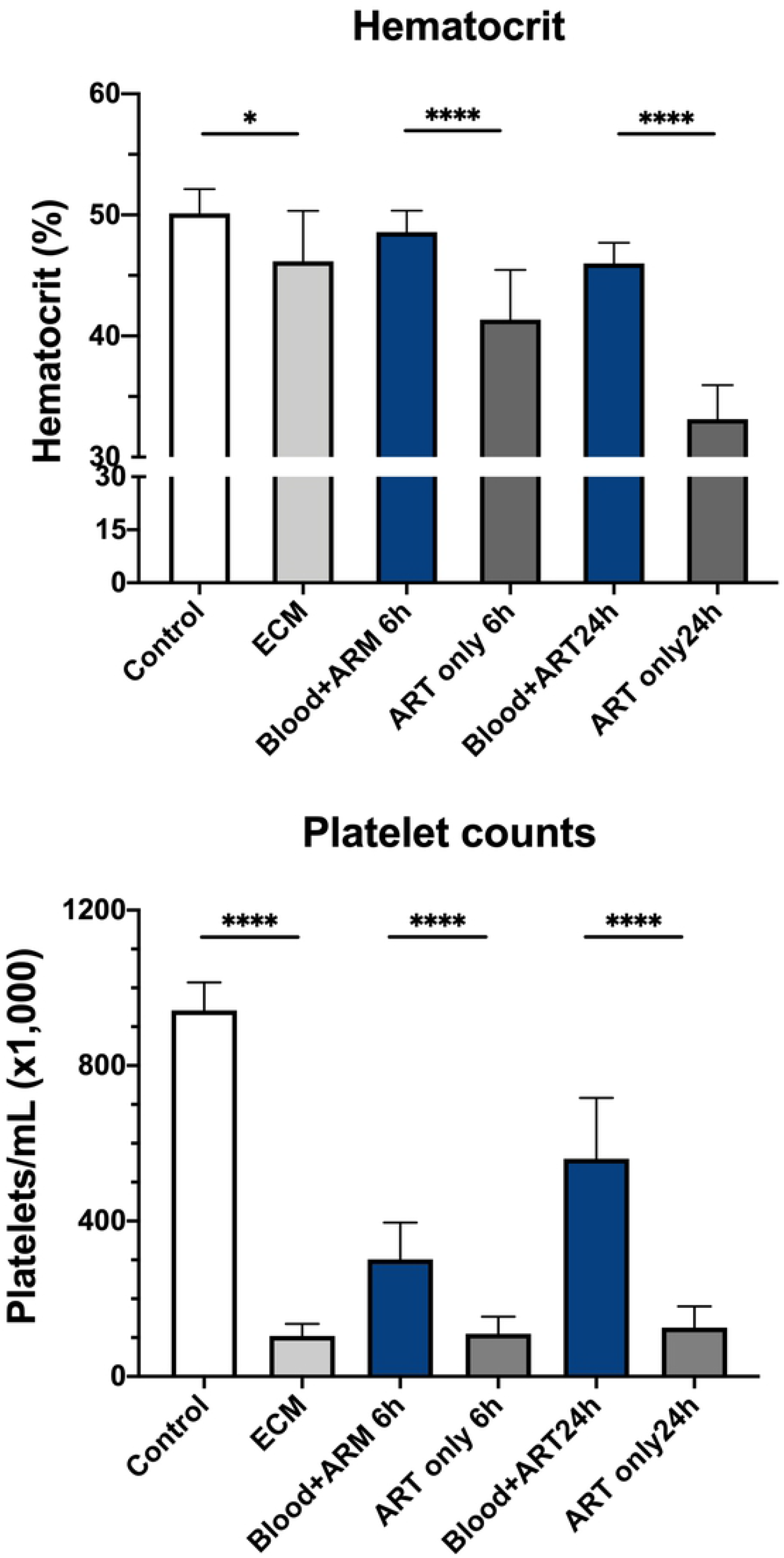
Improved survival after blood transfusion plus artemether compared to artemether alone and plasma plus artemether in mice with ECM. *Plasmodium-berghei* ANKA-infected mice showing signs of ECM and rectal temperatures between 31 and 36 °C on day 6 of infection were treated with either: *i*) artemether 20mg/kg given intraperitoneally (IP) alone (n = 13); *ii*) artemether IP plus 200 μL of whole blood given intravenously (IV) (n = 19) or; *iii*) artemether ip plus 200 μL of plasma IV (n = 11). Whole blood or plasma were given only once, together with first artemether dose, whereas artemether was given daily for a total of 5 days. Adjunctive therapy with whole blood resulted in marked improved survival (90%) compared to artemether alone (54%; p = 0.0004). On the other hand, associating plasma therapy with artemether resulted in worsened outcome, with only 18% survival. Three separate experiments were performed, and the results were combined. For statistical analysis, log-rank Mantell-Cox test was performed.

### Intravenous blood transfusion improves ECM-induced thrombocytopenia

The mean platelet count on day 6 post infection in mice with ECM was depleted by 90% (104,000 platelets per mL versus 942,000 of uninfected controls) (**Figure 1B**). Treatment with artemether alone had little effect on platelet counts at 6 hours and 24 hours. Intravenous blood transfusion as an adjuvant to artemether significantly hastened the recovery of platelets at 6 hours and at 24 hours (2.7-fold and 4.4-fold compared to ECM mice treated with artemether only, respectively). Intravenous blood transfusion recovered the platelet count to around 60% of that noticed in healthy controls, whereas artemether alone had no affect at any time point (13% of controls).

### Blood transfusion improves endothelial quiescence by restoring balance of angiopoietin-1 and 2

Plasma levels of ANG-1 and ANG-2 among four groups were assessed at different time points before and after treatment. With disease severity on day 6 post infection, plasma levels of endothelial-protective ANG-1 strongly declined (**Figure 2A**). Compared to uninfected healthy control, mean plasma levels of ANG-1 declined 84% in ECM mice. Treatment with artemether alone did not modify ANG-1 levels at 6 and 24 hours post treatment. Comparatively, at 24 hours mice receiving blood transfusion showed ANG-1 levels two times higher than artemether-only treated animals, indicating that transfusion induced a faster recovery of endothelial function in artemether-treated animals. On the other hand, the mean plasma levels of endothelial-inflammatory ANG-2 increased 52% in mice with ECM (**Figure 2B**). The levels of ANG-2 remained high 6 hours after either artemether or artemether plus blood transfusion. However, intravenous blood transfusion helped bring ANG-2 levels back to normal at 24 hours. As an additional measure, the ratio of ANG-2 to ANG-1 of ECM mice was found to be significantly different between artemether-only and artemether plus transfused blood groups at 24 hours (**Figure 2C**). No significant difference was noticed in the ratio of ANG-2 to ANG-1 between uninfected control and blood transfused groups. These data combined indicate that intravenous whole blood transfusion restores angiopoietin balance in artemether-treated ECM mice 24 hours after administration.

**Figure 2.**
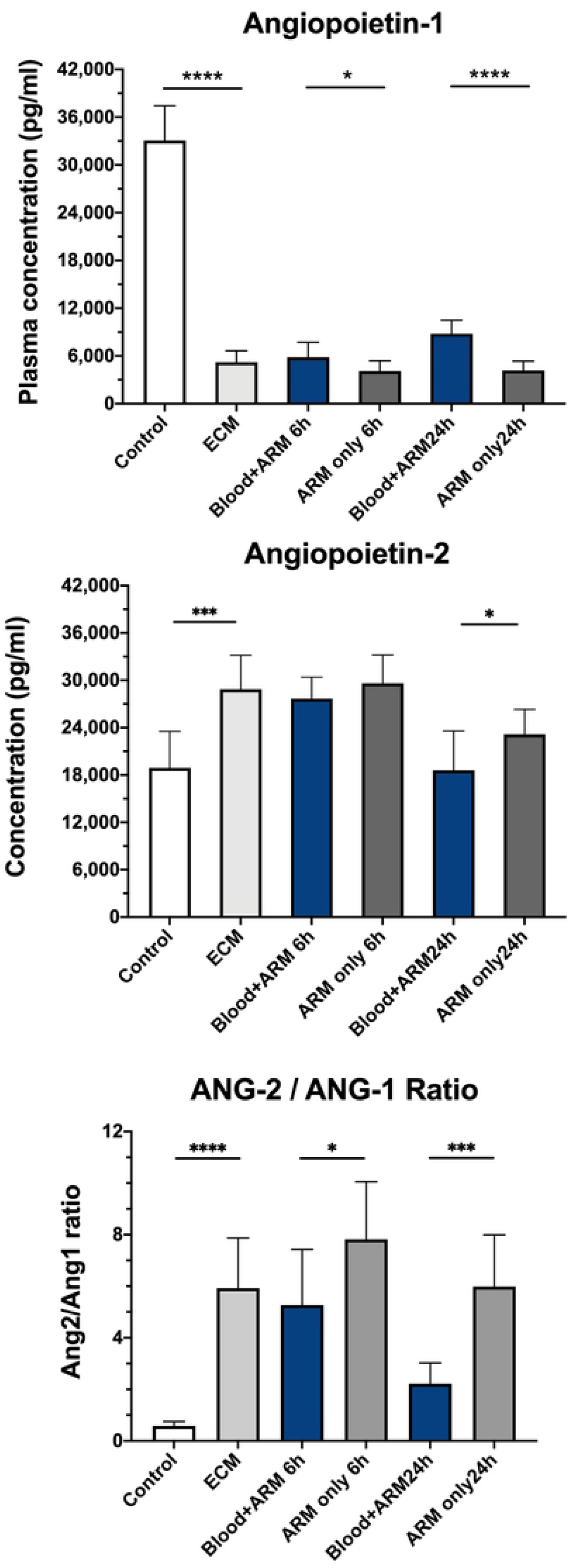
Effect of intravenous whole blood transfusion plus artemether on plasma angiopoietin-1 (ANG-1) and angiopoietin-2 (ANG-2) profiles in mice with ECM. Plasma levels of ANG among studied groups (healthy controls, ECM = experimental cerebral malaria, blood plus artemether [ARM] and artemether alone) were measured using ELISA (*n* = 10 mice per group). Mice with ECM showed depleted plasma ANG-1 levels (**A**) compared to uninfected controls (P < 0.0001). Treatment of ECM mice with artemether had no effect on ANG-1 levels, which remained very low. However, ANG-1 levels were slightly (mean 43%) but significantly higher in ECM mice that received whole blood transfusion compared to mice that received artemether alone at 6 hours (P = 0.0499) and at 24 hours whole blood transfusion led to a 2-fold increase in ANG-1 levels compared to artemether alone (P < 0.0001). (**B**) Plasma ANG-2 levels increased by 1.5 folds in ECM mice compared to normal controls (P = 0.0001). Treatment with artemether alone or artemether plus whole blood transfusion did not modify the increased ANG-2 levels 6 hours after treatment. At 24 hours, ECM mice that received whole blood transfusion showed plasma ANG-2 levels back to normal, whereas ECM mice that received artemether alone still showed increased levels (p = 0.0462). (**C**) The ratio of ANG-2 to ANG-1 was increased 10-fold in ECM mice compared to healthy controls (P < 0.0001). The levels kept increasing after 6 hours of artemether alone treatment but stabilized with artemether plus whole blood treatment (P = 0.0357). Within 24 hours, ANG-2 levels were still high in ECM mice that received artemether only, but were much lower in ECM mice that received artemether plus whole blood transfusion (P = 0.0002). Data are shown as mean ± standard deviation. For comparing groups at a given timepoint, Student’s t-test was performed.

### Intravenous blood transfusion prevents blood-brain barrier (BBB) breakdown

The improved survival of PbA-infected mice treated with whole blood in combination with artemether can be associated with preservation of the BBB, quantified by Evans blue extravasation into the brain parenchyma. Mice with ECM showed increased cerebrovascular Evans blue leakage, which was aggravated 6 hours post treatment with artemether alone (**Figure 3A**). Mice with ECM that received intravenous blood transfusion along with artemether showed no aggravation of vascular leakage, with Evans blue extravasation being much lower than that observed in mice that received artemether alone. Actually, these mice even showed some recovery of cerebrovascular integrity, as Evans blue leakage was significantly smaller than that observed in ECM mice prior to treatment. Also, after 24 hours of treatment, mice that received blood transfusion showed enhanced BBB integrity, comparable to the level of healthy controls, whereas animals treated with artemether alone still presented increased vascular leakage at this point.

**Figure 3:**
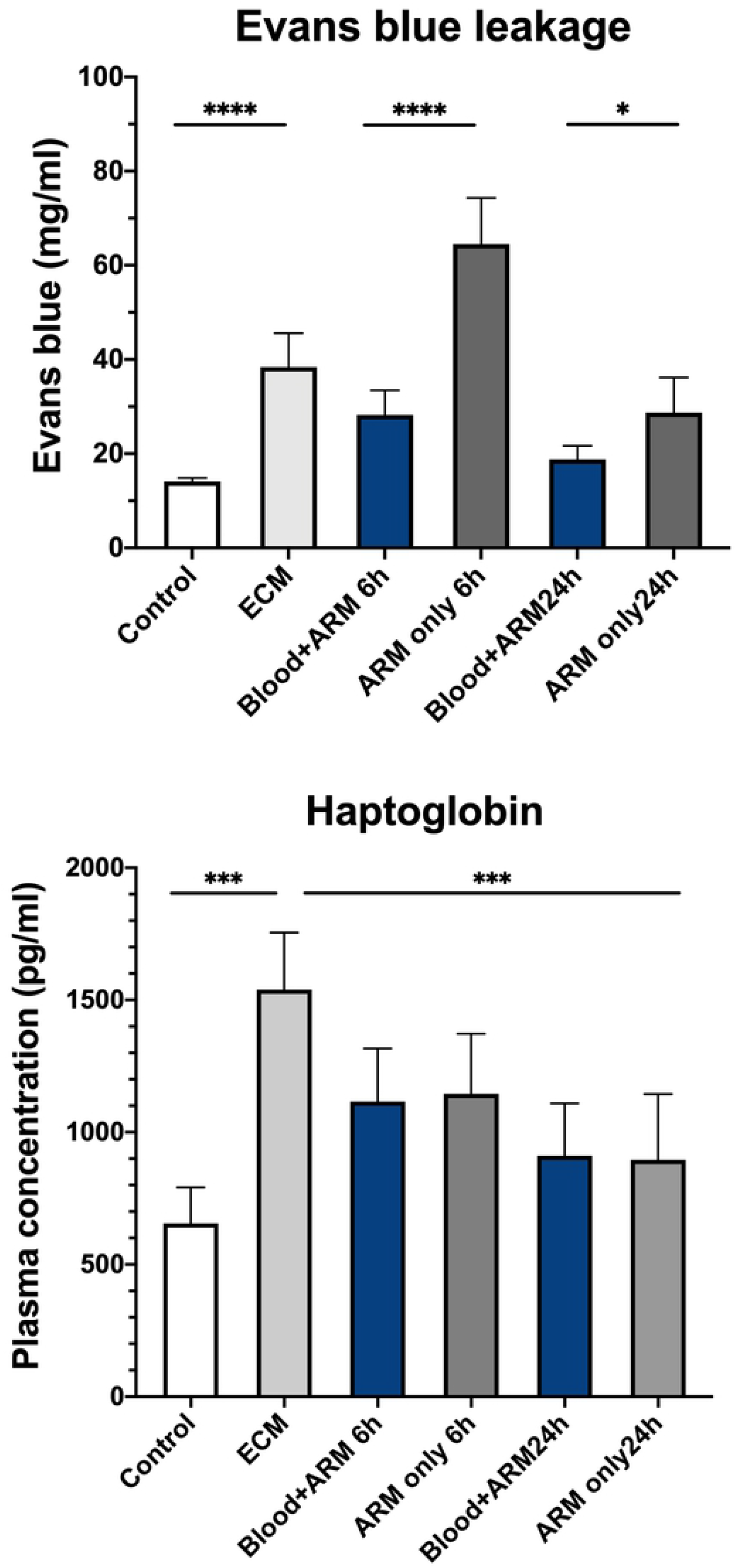
Blood-brain barrier (BBB) permeability and plasma levels of haptoglobin in mice with ECM after treatment with artemether with or without intravenous blood transfusion. **(A)** The BBB permeability (Evans blue assay) in mice with ECM was increased 2.7-fold in relation to healthy controls (P < 0.0001). BBB permeability kept increasing (to 4.6-fold) 6 hours after treatment with artemether (ARM) alone, but intravenous transfusion of whole blood prevented the further increase. Indeed, Evans blue leakage in the brain of transfused mice was less than half that observed in mice receiving artemether alone (p < 0.0001). At 24 hours, BBB leakage decreased but was still evident in mice treated with artemether alone, whereas it returned to normal levels in mice that received artemether plus blood transfusion (p = 0.0126). **(B)** Haptoglobin levels in whole plasma was assessed using ELISA. Mice with ECM showed a mean 135% increase in plasma haptoglobin levels compared to uninfected controls (P < 0.0001). The levels of haptoglobin showed a decline in both treatment groups at 6 and 24 hours. No difference between the treatments was noticed any time point. Data are shown as mean ± standard deviation. For comparing groups at a given timepoint, Student’s t-test was performed.

### Plasma haptoglobin levels after intravenous blood transfusion

Mice with ECM showed a mean 134% increase in plasma haptoglobin levels (1,539 ± 217 pg/mL) compared to healthy control (655 ± 136 pg/mL) (**Figure 3B**). Treatment with artemether, with or without intravenous blood transfusion, led to decreased haptoglobin levels at 6 hours (1,116 ± 201 pg/mL and 1,145 ± 227 pg/mL, respectively) and further at 24 hours (911 ± 198 pg/mL and 896 ± 248 pg/mL, respectively).

### Intravenous whole blood transfusion plus artemether improves survival of mice with ECM compared to artemether plus plasma or artemether alone

When PbA-infected mice present signs of ECM, if left untreated they will die, mostly within 24 hours. Treatment of mice with late-stage ECM with artemether led to a survival rate of 54% (**Figure 4**). Whole blood transfusion given together with artemether resulted in 90% survival, which represents a dramatic increase of 67% in survival compared to mice treated with artemether alone. Plasma transfusion (200 μL), on the other hand, worsened disease severity, resulting in very low survival rate (18%).

**Figure 4.**
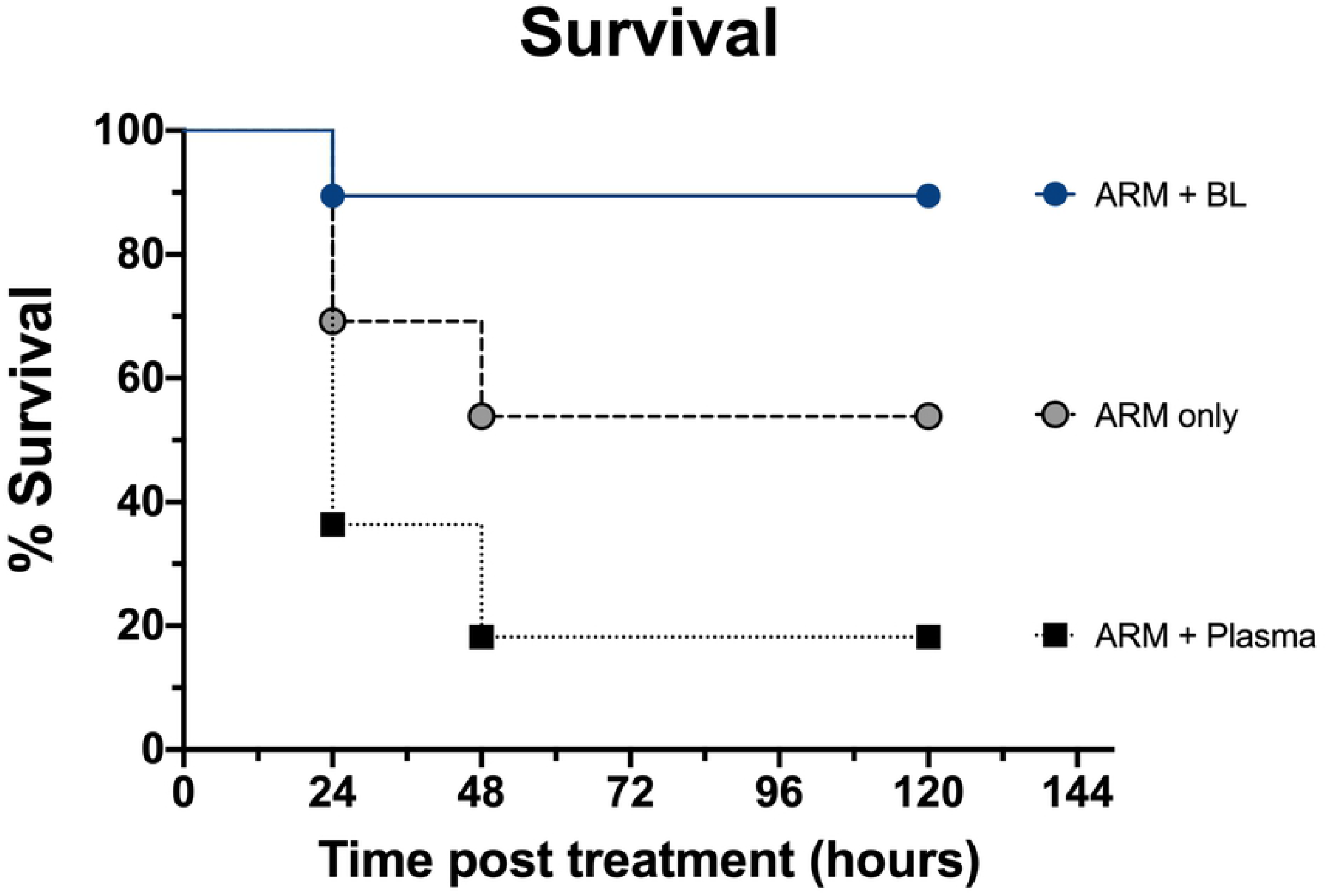
Effect of adjuvant intravenous blood transfusion at 6 and 24 hours on levels of hematocrit and platelet counts in mice with ECM. **(A) Hematocrit:** In ECM mice, pretreatment and pretransfusion hematocrit was 8% less than in healthy controls (mean; 46.16 % versus 50.12 %, P = 0.0390). Mice treated with artemether (ARM) alone showed further decrease in hematocrit after 6 and 24 hours (P < 0.0001). Intravenous blood transfusion plus ARM stabilized hematocrit levels at 6 and 24 hours (mean 48.8% and 46%, respectively). **(B) Platelet count:** ECM mice displayed around 90% reduction in platelet counts in relation to healthy controls (P < 0.0001). Intravenous blood transfusion led to partial recovery in platelet counts at 6 hours (2.9-fold; P < 0.0001) and at 24 hours (5.4-fold; P < 0.0001) compared to mice that received artemether alone. Data are shown as mean ± standard deviation. For comparing groups at a given timepoint, Student’s t-test was performed.

Overall, these findings suggest that whole blood, when transfused via intravenous route, improved survival and reduced disease severity by preserving endothelial components.

## Discussion

Clinical observational studies have suggested that intravenous whole blood transfusion may improve survival of children with cerebral malaria even in the absence of severe anemia. However, the mechanisms underlying the apparent protection of whole blood remain undefined. Here, we find that in a murine model cerebral malaria, intravenous whole blood restored the balance of angiopoietin 1 and 2, key factors that regulate endothelial inflammation and barrier integrity. Indeed, intravenous whole blood transfusion prevented the blood brain barrier breakdown that is a cardinal feature of experimental cerebral malaria, and improved survival. Notably, infusion of plasma instead of whole blood did not convey any protection and instead worsened survival. Thus, we must infer that red blood cells are a necessary component of endothelial-preserving property of whole blood.

The beneficial effects of blood transfusion include improvement in red blood cell deformability [26,27], maintenance of hematocrit strengthening the oxygen-carrying capacity of the blood, which in a setting of cerebral ischemia may be a critical advantage for the patient. Lower hematocrit also results in decreased vascular wall shear stress [28], with decreased eNOS activity, which together with the nitric oxide-scavenging action of free hemoglobin leads to worsened endothelial dysfunction. Under conditions of decreased oxygen saturation, fresh red blood cells may induce vasodilation of vessel strips by exporting NO bioactivity [29], release of ATP from RBCs also results in RBC-mediated hypoxic vasodilation [30,31]. Therefore, increasing hematocrit through whole blood transfusion should help restore endothelial function and improve tissue perfusion.

Prior work has demonstrated the therapeutic efficacy of whole blood transfusion in PbA-infected C57BL/6 mice with ECM, using intraperitoneal route for blood administration, resulting in 76% survival (compared to 51% survival in ECM mice receiving artemether only) [15]. The present study demonstrates that a single intrajugular transfusion of whole blood along with artemether resulted in a 90% survival of mice with ECM.

In the previous study, the intraperitoneal route of transfusion was chosen due to the difficulties of performing intravenous injection of large volumes in mice with ECM. However, the natural pathway for blood transfusion is the intravenous route, and a solution was achieved by means of intrajugular injection. This strategy resulted in a stronger increase in survival, and this effect was related to a more effective action on measured parameters of hematological and vascular health. Indeed, the findings in the present study independently confirm the protective effects of intraperitoneal whole blood transfusion, and now extend those findings to the more clinically relevant intravenous route of administration. Interestingly, intravenous transfusion appeared to have a greater effect on hematocrit, platelet counts, ANG-1 and ANG-2 levels, blood-brain barrier integrity, and survival compared to intraperitoneal transfusion.

Endothelial cell system activation and dysfunction are hallmarks of malaria pathogenesis [32–34]. Endothelial protein markers such as angiopoietins have been found to play a critical physiological role in maintenance of vascular integrity. It has been previously shown that the levels of the endothelial cell quiescence promoter ANG-1 are depleted, whereas the levels of endothelial cell activation/inflammation promoter ANG-2 are elevated, in mice with ECM [15]. It is noteworthy that blood transfusion given intraperitoneally had no effect on restoring ANG-1 levels, but when given intravenously a timid but significant increase in ANG-1 levels was observed after 6 hours, and a more robust increase was seen at 24 hours. And while ANG-2 levels were equally restored by either intraperitoneal of intravascular transfusion, the better effect on ANG-1 levels resulted also in a much better ANG-2/ANG-1 ratio by intravascular transfusion. Since ANG-1 and ANG-2 compete for binding to Tie2, with opposite effects, this improved profile achieved by intravascular transfusion likely results in a faster recovery of endothelial function. Indeed, reversing ANG-2/1 imbalance promotes endothelial cell survival, stabilizes endothelial interactions with supporting cells and limits vascular permeability [35–37]. ANG-1 stabilizes the BBB while ANG-2 weakens the pericyte-endothelial cell interaction, resulting in BBB disruption [34,38]. Since severity of cerebral malaria is related to vascular dysfunction [39] and much of the mortality occurs in the first 24 hours after antimalarial treatment, this faster recovery of endothelial function may be a critical factor explaining the substantial increase in survival in these animals.

A faster recovery of endothelial function may help to explain a more rapid reversal of BBB breakdown. The overall profile of Evans blue leakage observed after treatment of ECM mice with artemether or artemether plus intravenous blood transfusion was similar to that observed in the previous study using intraperitoneal transfusion [15]. However, ECM mice receiving intravenous transfusion showed a more profound recovery of BBB integrity, both at 6 and 24 hours after treatment. These findings are in line with a better profile of plasma ANG-1 and ANG-2/ANG-1 levels and with a better survival outcome. In the case of haptoglobin levels, the effects of intravenous transfusion were not different from those observed with intraperitoneal transfusion [15].

Malaria is a true hematological infectious disease, heavy parasite burden and hemolysis of parasitized and non-parasitized red blood cells may result in rapid decline of hematocrit [40,41]. Intravenous transfusion again resulted in a better hematological outcome compared to intraperitoneal transfusion. Whereas the latter only partially prevented the marked fall of hematocrit following artemether treatment observed in non-transfused mice [15], intravenous transfusion resulted in a more consistent maintenance of hematocrit. This can be in part explained by the fact that not all red blood cells transfused by intraperitoneal route actually reach the circulation [24]. But there is also indication that blood transfusion does not lead only to a passive recovery of circulating red blood cell numbers. The magnitude of hematocrit recovery at 24 hours in transfused compared to non-transfused mice was greater than it would be expected by the amount of red blood cells transfused. Therefore, whole blood transfusion may either induce an active response of the recipient mice, enhancing bone marrow production of new red blood cells, or transfusion has a protective effect on the hemolytic effect of artemether. The actual mechanisms explaining these findings still need to be shown.

In any case, the maintenance of higher hematocrit has beneficial effects such as increasing shear stress [28], managing acid-base balance [13,42], improving red blood cell deformability, reducing microvascular obstruction, all of which help improving brain tissue oxygen delivery [43,44] [45].

Another one of the more well-known hematologic changes observed in patients with malaria is thrombocytopenia [46]. *Plasmodium falciparum* infected patients with a platelet count below 20 × 10^9^/L were five times more likely to die than patients with a higher platelet count [47]. Intraperitoneal blood transfusion partially restored platelet counts in ECM mice [15]. But, again, the effect of intravenous blood transfusion was stronger, leading to a faster recovery of platelet counts and again the magnitude of the recovery was greater than it would be expected by a passive transfer of platelets contained in the transfused blood. Because platelets have been implicated with the pathogenesis of ECM, in processes such as coagulopathy [48], endothelial activation [32], cytoadherence [49] and auto-agglutination [50], restoring fresh platelets in a timely manner may be helpful in preventing mortality.

Finally, this study confirmed that, while whole blood transfusion is an intervention with high efficacy in preventing death by ECM, healthy plasma transfusion has the opposite effect, leading to increased mortality. This is probably related to increased circulating volume without increased oxygen carrying capacity which may worsen cardiovascular performance exacerbate pulmonary edema [51,52].

Collectively our data suggest that intravenous blood transfusion reversed anemia and thrombocytopenia, facilitated restoration of endothelial quiescence, improved vascular stability, leading to improved survival in experimental CM. These findings deserve more scrutiny to better understand the mechanisms behind the benefit achieved and provide additional support for translational studies.

## Author contributions

SG conducted the experiments, analyzed the data and wrote the article. HCA was involved with study conception and, with CTDR, helped with data analysis and interpretation. LJMC conceived and was overall responsible for planning and supervising the study, data analysis and interpretation, and wrote the manuscript. All authors reviewed the manuscript.

## Acknowledgements

We thank Dr Tatiana Maron Gutierrez and Dr Helena D’Anunciação de Oliveira (IOC-Fiocruz) for training SG in the technique of blood transfusion via the jugular vein, and Cleber Hooper (ICTB-Fiocruz) for allowing access to the hematological analyses facilities and running the mouse blood samples. This work was supported by the ‘Universal’, “Productivity” and ‘INCT-NIM’ grants from the Brazilian National Research Council (CNPq) (LJMC, CTDR), by the ‘Cientista do Nosso Estado’ grant from the Rio de Janeiro Research Funding Agency (Faperj) (LJMC, CTDR), by the Oswaldo Cruz Institute and in part by the Intramural Research Program of the NIH (HCA). SG received a fellowship from CAPES.

